# Actin filament oxidation by MICAL1 suppresses protections from cofilin-induced disassembly

**DOI:** 10.1101/2020.02.26.966614

**Authors:** Hugo Wioland, Stéphane Frémont, Bérengère Guichard, Arnaud Echard, Antoine Jégou, Guillaume Romet-Lemonne

**Affiliations:** Université de Paris, CNRS, Institut Jacques Monod, 75013 Paris, France; Membrane Traffic and Cell Division Lab, Institut Pasteur, UMR3691, CNRS, F-75015 Paris, France

## Abstract

Proteins of the ADF/cofilin family play a central role in the disassembly of actin filaments, and their activity must be tightly regulated in cells. Recently, the oxidation of actin filaments by the enzyme MICAL1 was found to amplify the severing action of cofilin through unclear mechanisms. Two essential factors normally prevent filament disassembly: the inactivation of cofilin by phosphorylation, and the protection of filaments by tropomyosins, but whether actin oxidation might interfere with these safeguard mechanisms is unknown. Using single filament experiments *in vitro*, we found that actin filament oxidation by MICAL1 increases, by several orders of magnitude, both cofilin binding and severing rates, explaining the dramatic synergy between oxidation and cofilin for filament disassembly. Remarkably, we found that actin oxidation bypasses the need for cofilin activation by dephosphorylation. Indeed, non-activated, phosphomimetic S3D-cofilin binds and severs oxidized actin filaments rapidly, in conditions where non-oxidized filaments are unaffected. Finally, tropomyosin Tpm1.8 loses its ability to protect filaments from cofilin severing activity when actin is oxidized by MICAL1. Together, our results show that MICAL1-induced oxidation of actin filaments suppresses their physiological protection from the action of cofilin. We propose that in cells, direct post-translational modification of actin filaments by oxidation is a way to trigger their severing, in spite of being decorated by tropomyosin, and without requiring the activation of cofilin.

## INTRODUCTION

The controlled assembly and disassembly of actin filaments (F-actin) is essential for numerous fundamental cellular functions. On one hand, a myriad of actin binding proteins (ABPs) have been identified and implicated in filament polymerization and spatial organization, force generation, and actin disassembly. On the other hand, actin post-translational modifications (PTMs) recently emerged as key players of actin dynamics. But the role and consequences of PTMs at the molecular level often remain mysterious (Varland et al., 2019).

Actin oxidation is one of such important PTM. The family of oxidoreductases MICAL, in the presence of the coenzyme NADPH and O_2_, has been identified to oxidize actin, its main target (Bai et al., 2020; Frémont et al., 2017a, 2017b; Grintsevich et al., 2017; Hung et al., 2010, 2011). Here, we focus on MICAL1, one of the three members of the MICAL family in humans. MICAL1 specifically targets F-actin to oxidize actin Met44 and Met47 (Hung et al., 2011)(Bai et al., 2020; Frémont et al., 2017a; Grintsevich et al., 2017); (Hung et al., 2011)(Bai et al., 2020; Frémont et al., 2017a; Grintsevich et al., 2017). These two residues are part of the “D-loop”, a region of the actin monomer involved in bridging interstrand and intrastrand actin subunits and thus important for actin filament stability (Chou and Pollard, 2019; Dominguez and Holmes, 2011; Grintsevich et al., 2017). As a consequence, MICAL1-mediated oxidation strongly accelerates actin filament depolymerization *in vitro* (Bai et al., 2020; Frémont et al., 2017a; Grintsevich et al., 2017). Oxidation by MICAL1 is selectively reversed by a family of methionine sulfoxide reductases (MsrB) (Bai et al., 2020; Hung et al., 2013; Lee et al., 2013). MsrB specifically targets oxidized monomeric actin (G-actin) and reduces Met44 and Met47, allowing for the repolymerization of actin filaments (Bai et al., 2020; Hung et al., 2013; Lee et al., 2013).

Actin oxidation by MICAL1 plays important roles to regulate actin assembly in muscle organization, axon guidance and drosophila bristle development. More recently, it has been implicated at the end of cell division, to clear actin filaments from the intercellular bridge, a crucial step for successful abscission (Frémont et al., 2017a). The activity of MICAL1 is balanced with that of MsrB2 which slows down the disassembly of actin structures during cytokinesis (Bai et al., 2020; Hung et al., 2013; Lee et al., 2013).

*In vitro*, oxidation by MICAL1 accelerates filament depolymerization, but, in contrast to initial reports (Hung et al., 2011), it does not induce filament fragmentation (Frémont et al., 2017a)(Grintsevich et al., 2016); (Frémont et al., 2017a). In a cellular context, MICAL1 is thus most likely cooperating with other molecular partners to efficiently disassemble actin filaments.

A central regulator of actin filament disassembly is the ubiquitous family of proteins ADF/cofilin (Elam et al., 2013; Hild et al., 2014; Kanellos and Frame, 2016). ADF/cofilin is commonly known to fragment actin filaments and can induce their depolymerization from both filament ends (Wioland et al., 2017, 2019a). Numerous studies described the action of ADF/cofilin at the molecular scale: it binds cooperatively to filaments, forming domains that locally modify actin filament conformation (Huehn et al., 2018, 2020; McGough et al., 1997; Prochniewicz et al., 2005; Wioland et al., 2019b) and induces severing at ADF/cofilin domain boundaries (Gressin et al., 2015; Suarez et al., 2011; Wioland et al., 2017). Nevertheless, on its own, ADF/cofilin cannot account for the very rapid turn-over of filaments observed in cells. It can be assisted by various other ABPs, such as Cyclase Associated Protein which accelerates the pointed end depolymerization of cofilin-decorated filaments (Kotila et al., 2019; Shekhar et al., 2019).

Interestingly, Grintsevich *et al*. recently identified a synergy between dMICAL from Drosophila and the human isoform cofilin-1: when oxidized by dMICAL, filaments are rapidly fragmented by cofilin-1 (Grintsevich et al., 2016). The number of cofilin domain boundaries, where severing can occur, results from the balance between the formation of new domains and their growth rate, and it is unclear how these two rates are affected by filament oxidation. In addition, the impact of filament fragilization by oxidation (Grintsevich et al., 2016) on the severing rate per cofilin-1 domain is unknown.

*In vivo*, cofilin-1 is a very abundant protein, reaching tens of µM (Bekker-Jensen et al., 2017). Its activity is regulated by various factors. A prevalent factor is cofilin-1 Ser3 phosphorylation by LIM kinases (Arber et al., 1998; Kanellos and Frame, 2016). This phosphorylation strongly and persistently inhibits cofilin-1 as it reduces its affinity for actin filaments and slows down subsequent severing (Elam et al., 2017). Additionally, cofilin-1 activity can be restrained by tropomyosins (Tpm) (Brayford et al., 2016; Christensen et al., 2017; Gateva et al., 2017; Jansen and Goode, 2019). Tpm is a long dimeric protein which binds and saturates most actin filaments in cells (Meiring et al., 2018). Tpm isoforms regulate specifically the recruitment of other ABPs and most Tpm isoforms prevent cofilin-1 binding (Gateva et al., 2017; Gunning and Hardeman, 2017; Jansen and Goode, 2019; Manstein et al., 2019). The interplay between Tpm decoration and post-translational modifications of actin has never been tested.

Here, we used single filament microfluidics experiments to monitor and quantify the action of cofilin-1 (Figure 1) and phosphomimetic S3D-cofilin-1 (Figure 2) on oxidized filaments compared to standard, non-oxidized filaments. We next measured whether MICAL1 can oxidize Tpm1.8-saturated filaments (Figure 3). Finally, we quantified the extent to which Tpm1.8 protects non-oxidized and oxidized filaments from cofilin-1-induced severing (Figure 4). The results lead us to conclude that the oxidation of filaments by MICAL1 is a way to trigger their severing, in spite of their being decorated by tropomyosin, and without requiring the activation of cofilin-1 (Figure 5).

**FIG 1 :**
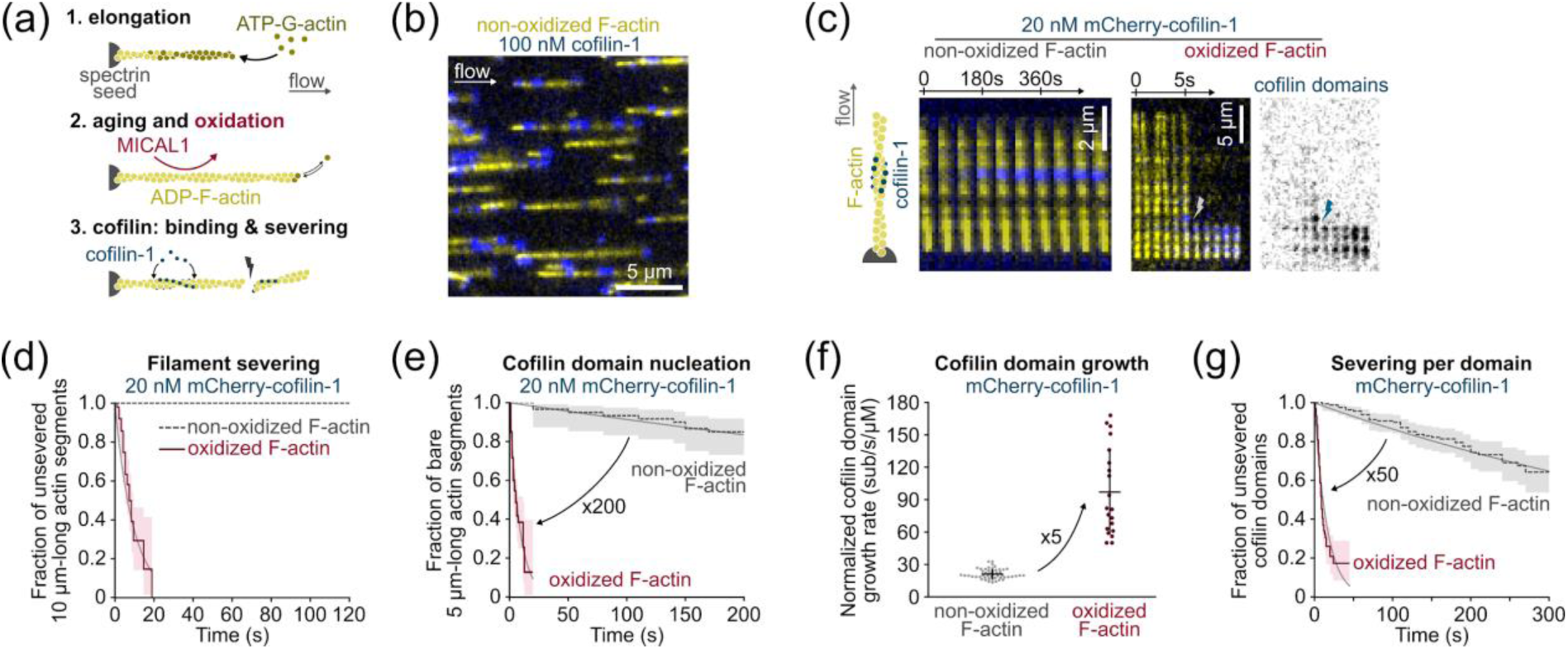
Cofilin at low concentration quickly binds and severs oxidized actin filament. (a) Three steps of a typical experiment (see also Methods). Filaments are polymerized with 0.6-1 µM ATP-G-actin and aged for 15 min with ATP-G-actin at critical concentration (0.1 µM) to maintain the filament length. This solution is supplemented with MICAL1 and NADPH to oxidize filaments. Tpm can also be added at this step to fully decorate filaments (Figure 3 and 4). (b) Fraction (1/17th) of a typical field of view. In a microfluidic chamber, actin filaments (yellow) are anchored by their pointed ends and align with the flow. (c) Kymograph showing the assembly of cofilin-1 domains (blue) and subsequent disassembly of actin filaments (yellow). Right: the kymograph of an oxidized filament shows filament severing at 6s; the first cofilin domain nucleation at 2s; growth of 4 individual domains as their fluorescence intensity increases; and severing (lightning symbol) at the top domain 2s after its nucleation. (d) Global measurement of the severing of filaments exposed to 20 nM mCherry-cofilin-1 from time t=0 onwards. N = 40 and 50 non-oxidized and oxidized actin filaments respectively. P-value < 0.001. (e) Nucleation of the first cofilin domain onto 5 µm-long actin segments. Filaments are exposed to 20 nM mCherry-cofilin-1 from time t=0 onwards. N = 60 filaments for the two conditions. P-value < 0.001. (f) Growth rate of single cofilin domains, normalized by the cofilin concentration. N = 50 and 20 domains on non-oxidized and oxidized filaments respectively. Measurements were obtained using 20 and 100 nM cofilin (non-oxidized actin) and 10 nM cofilin (oxidized actin). Note that this normalized growth rate does not depend on the cofilin concentration (Supp Fig 1a). Bars: mean and S.D. P-value < 0.001. (g) Filament severing rate at single cofilin domains. Time t=0 is defined for every domain as the frame on which they nucleate. N = 163 and 203 cofilin domains on non-oxidized and oxidized actin respectively. Measurements were performed at 100 nM cofilin (non-oxidized actin) and 10, 20 or 30 nM cofilin (oxidized actin). Since the severing rate does not depend on the cofilin concentration (Supp Fig 1b and (Wioland et al., 2017, 2019a)) data for 10, 20 and 30 nM cofilin on oxidized filaments were pooled. Measurements on non-oxidized filaments were done at 100 nM mCherry-cofilin-1. P-value < 0.001. (d, e, g) Thin grey lines are single exponential fits. 95% confidence intervals are shown as shaded surfaces.

**FIG 2 :**
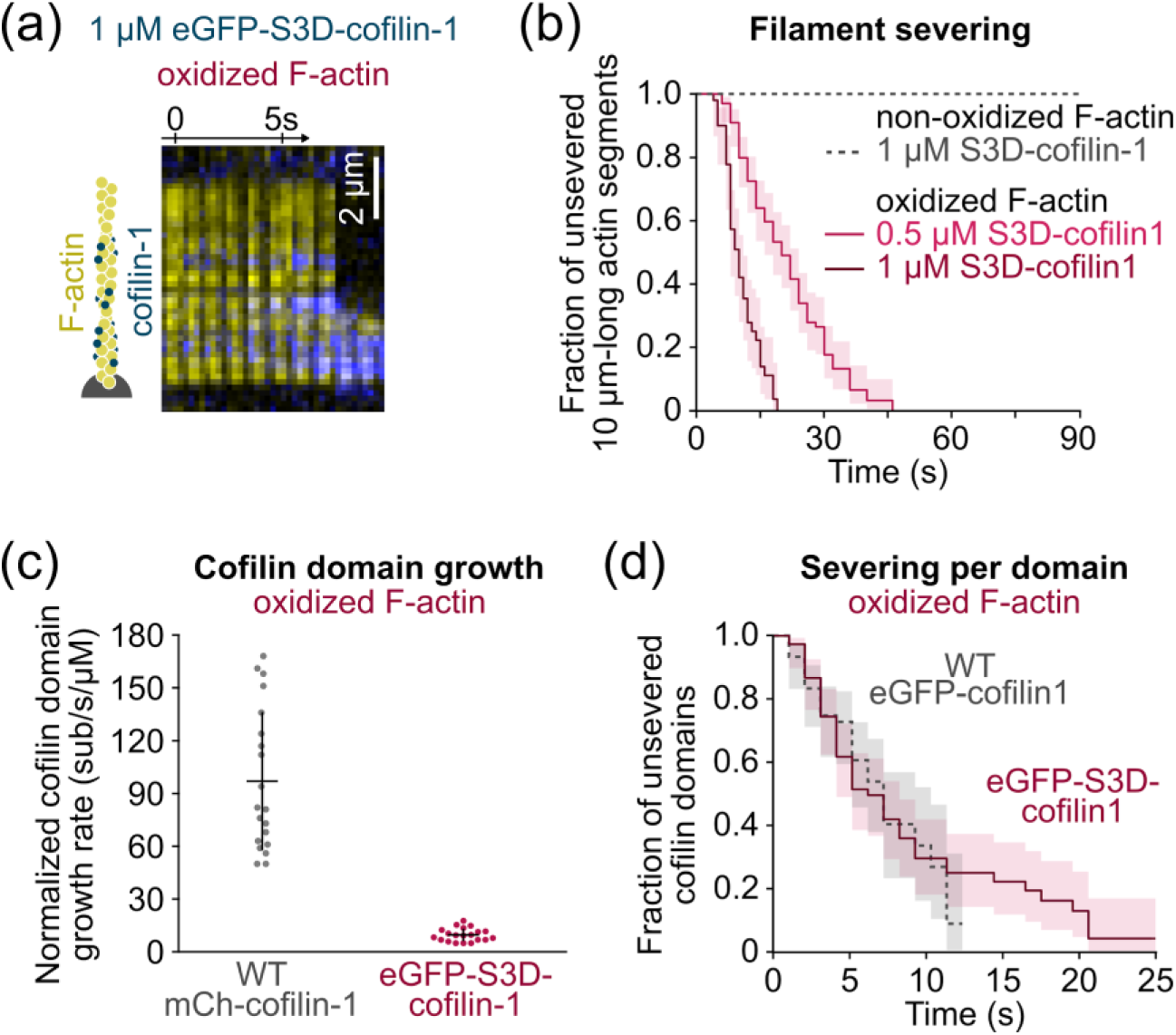
Phospho-mimic (S3D) cofilin efficiently binds and severs oxidized F-actin. (a) Time-lapse images of eGFP-S3D-cofilin-1 (blue) binding and severing of an oxidized actin filament (yellow). (b) Global quantification of the severing events on oxidized and non-oxidized filaments exposed to unlabelled S3D-cofilin1 from time t=0 onwards. From top to bottom, N = 50, 50, 100 filaments. (c) Growth rate of single cofilin domains, normalized by the cofilin concentration. N = 20 domains (1 and 2 experiments). The growth rate was measured at 10 nM WT cofilin, and 500 nM and 1 µM S3D-cofilin-1 (Supp Fig 1c). Bars: mean and S.D. P-value < 0.001. (d) Filament severing rate at single cofilin domains. Time t=0 is defined for every domain as the frame on which they nucleate. N = 60 and 75 cofilin domains on non-oxidized and oxidized actin respectively. The severing dynamic was measured at 10 nM WT eGFP-cofilin-1 and 1 µM eGFP-S3D-cofilin-1. P-value = 0.98. (b, d) Thin grey lines are single exponential fits. 95% confidence intervals are shown as shaded surfaces.

**FIG 3 :**
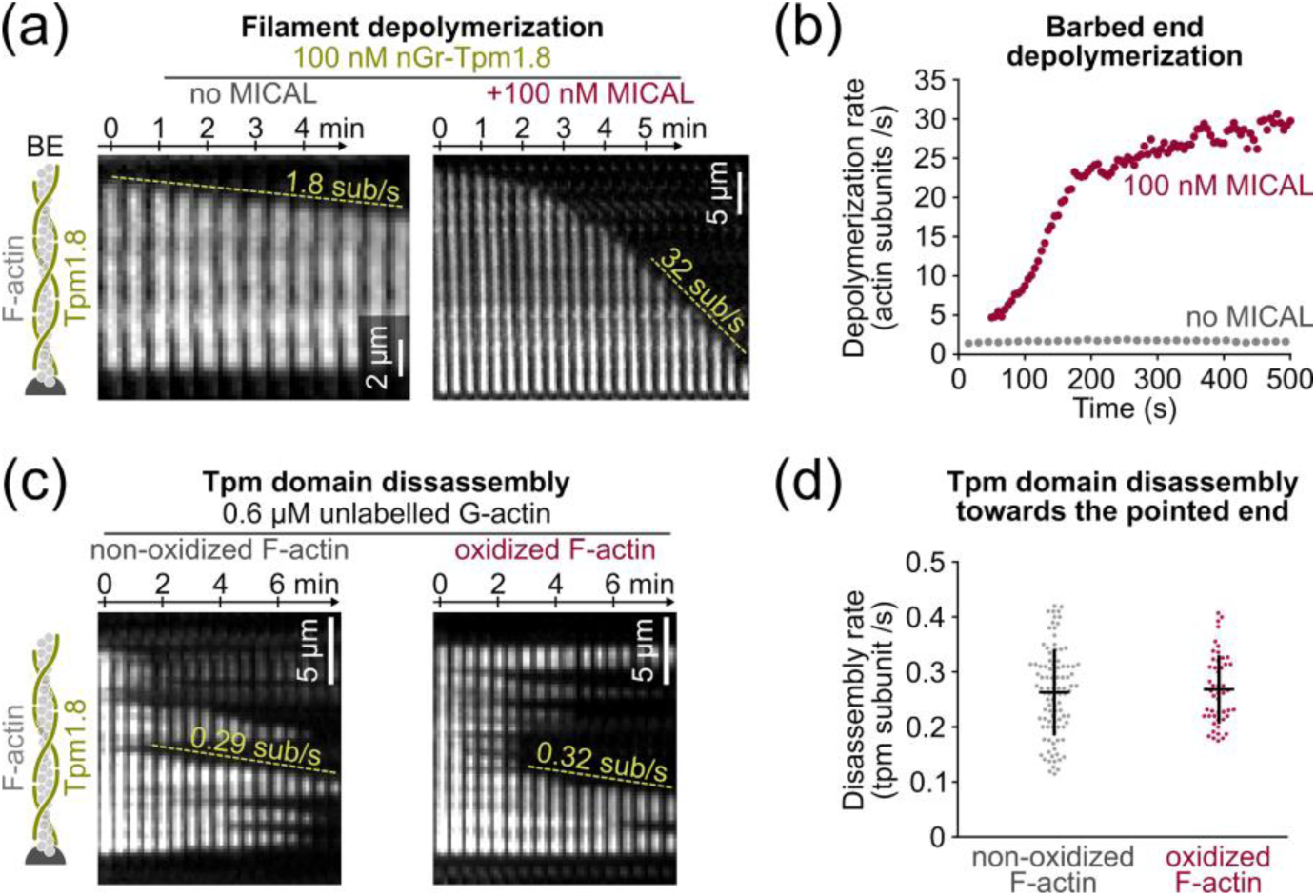
MICAL1 can oxidize Tpm-saturated filaments and does not affect Tpm unbinding. (a) Time-lapse images of Tpm-saturated actin filament depolymerizing from their barbed end (BE), with or without MICAL. Depolymerization is initiated at time t=0 from a non-oxidized actin filament. Right: 100 nM MICAL1 and 12 µM NADPH are injected from t=0 onwards. The increase in the depolymerization rate reflects the oxidation of actin with time. Tpm is labeled with Neon Green, and actin is unlabeled. (b) Barbed end depolymerization rate as a function of the exposure time to either 100 nM nGr-Tpm1.8 alone (“no MICAL”), or supplemented with 100 nM MICAL1 and 12 µM NADPH. N = 25 and 28 filaments, without and with MICAL1 respectively. (c) Time-lapse images showing the disassembly of Tpm domains. Filaments were saturated with 100 nM nGr-Tpm1.8 before the experiment and Tpm was removed from solution at time t=0 (the solution contains 0.6 µM G-actin to prevent filament depolymerization). (d) Disassembly rate of single Tpm domains. As domains disassemble faster towards the pointed end, only the dissociation towards the pointed end anchor (actin-spectrin seed) was taken into account to obtain more accurate results. Left to right: N = 100 and 50 domains. Bars: mean and S.D. P-value = 0.68.

**FIG 4 :**
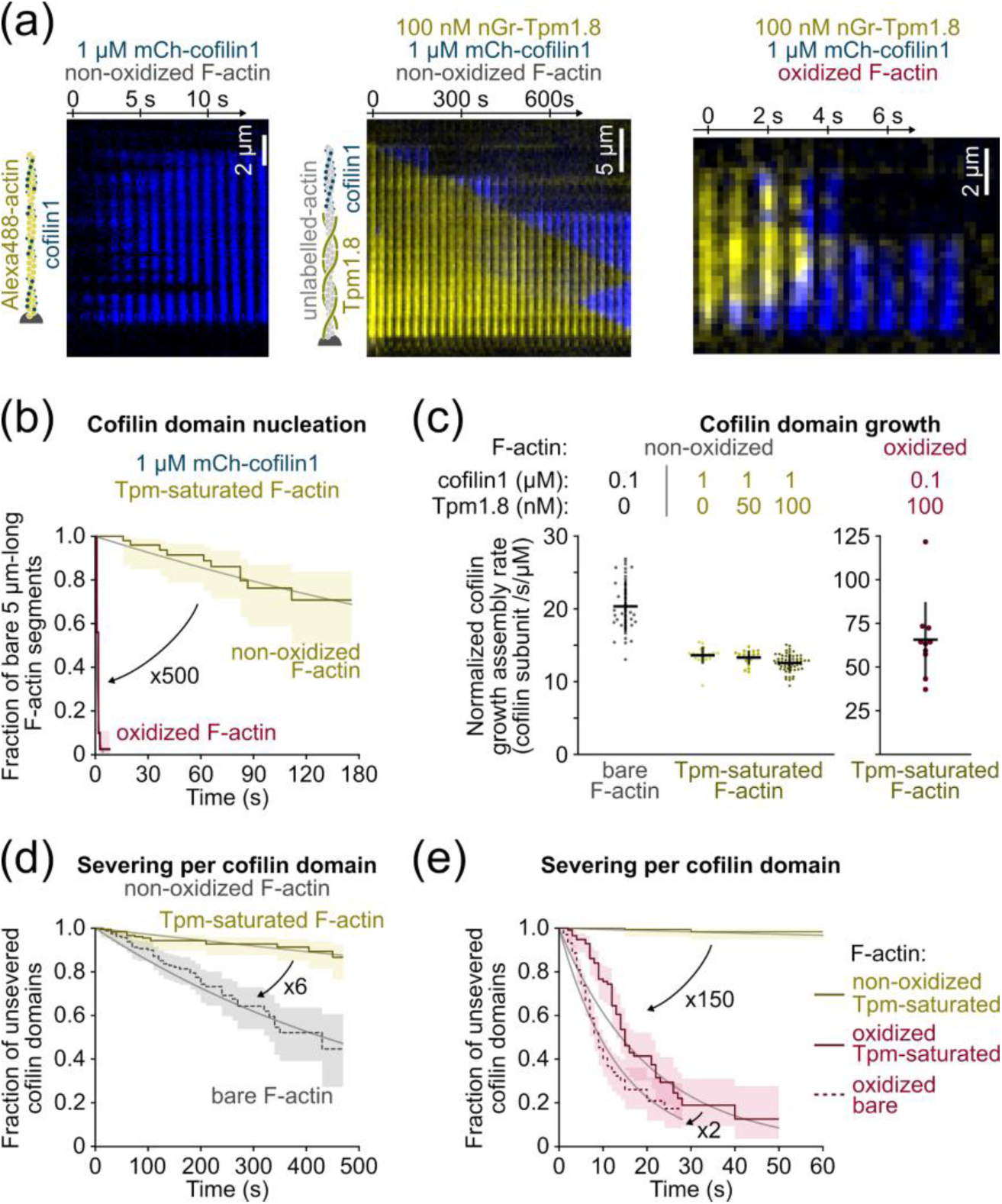
Tpm protects non-oxidized but not oxidized F-actin from cofilin-induced severing. (a) Time-lapse images of filaments, oxidized or not, bare or Tpm-saturated, exposed to 1 µM mCherry-cofilin-1, from time t=0 onwards. (b) Nucleation of the first cofilin domain onto 5 µm-long Tpm-saturated actin filaments. Filaments are exposed to 1 µM mcherry-cofilin from time t=0 onwards. N = 50 filaments for each condition. (c) Growth rate of single cofilin domains, normalized by the cofilin concentration. Filaments saturated by Tpm during aging (Figure 1a) were then exposed to cofilin and 0-100 nM Tpm. Bare filaments exposed to cofilin in the absence of Tpm are shown as a reference (left). Note the scale difference for oxidized filaments (right). From left to right, N = 40, 20, 20, 60, 10 domains. Bars: Mean and S.D. (d-e) Filament severing rate at single cofilin domains. Time t=0 is defined for every domain as the frame on which they nucleate. N = 164, 180, 116 domains for bare actin (100 nM cofilin), non-oxidized Tpm-saturated (1 µM cofilin) and oxidized Tpm-saturated filaments (100 nM cofilin), respectively. (b, d, e) Thin grey lines are single exponential fits. 95% confidence intervals are shown as shaded surfaces.

**FIG 5 :**
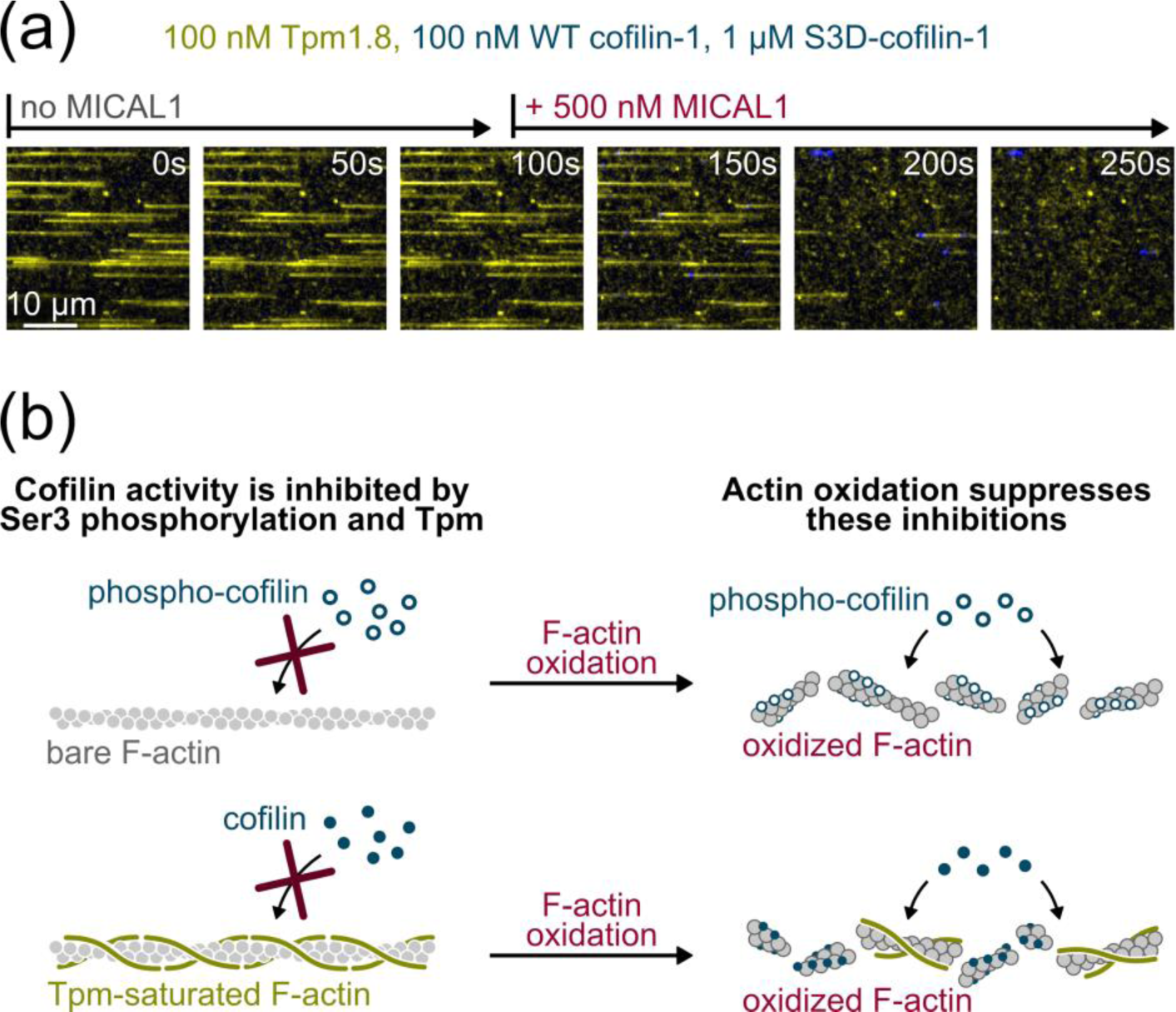
Actin filament oxidation removes their protections from cofilin-induced disassembly. (a) Time-lapse of actin filaments constantly exposed to Tpm1.8 (yellow), WT cofilin-1 (blue) and S3D-cofilin-1, and to MICAL1 from time t=100s onwards. While Tpm prevents cofilin from binding to non-oxidized filaments, MICAL1 rapidly oxidizes actin, allowing cofilin to bind and sever filaments. See also Supp. Movie. (b) Summary of the results. Different mechanisms can simultaneously down regulate the activity of cofilin (here cofilin phosphorylation and protection by Tpm). Actin oxidation is a rapid means to cancel these protections without, for example, activating a large concentration of cofilin.

## RESULTS

### MICAL1-induced actin oxidation enhances both cofilin-1 binding and subsequent severing

In order to dissect the action of cofilin-1 on oxidized and non-oxidized filaments, we used a microfluidic setup previously developed in the lab (Jégou et al., 2011; Wioland et al., 2019c). Briefly, filaments were polymerized from surface-anchored Spectrin-actin seeds (Figure 1a). Filaments were then aged for 15 minutes to become fully ADP-actin filaments. During that time, the actin filaments were also exposed to MICAL1 to be oxidized, or not (Frémont et al., 2017a). Finally, we exposed them to mCherry-cofilin-1. All experiments were carried out at pH 7.4 and at room temperature, with alpha-skeletal actin (rabbit), recombinant human cofilin-1 and S3D-cofilin-1, recombinant human pseudo-N-acetylated Tpm1.8, and the recombinant active domain of human MICAL1 (see Methods for details).

We first assessed the impact of actin oxidation by MICAL1 on the activity of wild type (WT, i.e. non-phosphorylated) cofilin-1. At a low concentration of cofilin-1 (20 nM), we observed no severing on non-oxidized filaments after 120s. In contrast, oxidized filaments quickly severed and none remained intact after 20s (figure 1 c, d), in agreement with previous work (Grintsevich et al., 2016). This accelerated disassembly could be due to either altered cofilin-1 binding or subsequent accelerated severing. We thus independently measured cofilin-1 domain nucleation rate, domain growth rate, and filament severing rate per cofilin-1 domain.

The nucleation rate was estimated by continuously exposing filaments to 20 nM mCherry-cofilin-1 and quantifying, over time, the fraction of 5 µm-long actin segments with no visible cofilin-1 domain. While cofilin-1 domains appeared very slowly on non-oxidized filaments (less than 20% had a cofilin-1 domain after 200s), nearly all oxidized segments presented one or more domain after 20s. Overall, actin oxidation increases by 200-fold the nucleation rate of cofilin-1 domains (from 3×10^−7^ s/sub to 7×10^−5^, figure 1e).

Similarly, we compared the growth rate of single cofilin-1 domains onto non-oxidized and oxidized actin filaments. To do so, we used different concentrations of cofilin-1, high enough for domains to steadily appear and grow, and low enough to limit nucleation and be able to track single domains. While mCherry-cofilin-1 domains grow by ∼20 sub/s/µM on non-oxidized filaments, this rate increased 5-fold, to ∼100 sub/s/µM, on oxidized filaments (Figure 1f, Supp Fig 1). Both nucleation and growth quantifications show that actin oxidation by MICAL1 largely increases the affinity of cofilin-1 for the filament. As a result, numerous domains are generated, in seconds (figure 1c).

The global cofilin-induced filament severing dynamic is the product of the number of cofilin-1 domains (at the boundary of which severing occurs, Figure 1c) and the severing rate per domain. To quantify the latter, we measured the fraction of unsevered cofilin-1 domains, from their nucleation to severing or loss (due for example to the severing at another domain). Fits of the survival fraction of domains with a single exponential showed that severing per domain was 50-fold faster on oxidized filaments (from 2×10^−3^ /s to 7×10^−2^ /s, Figure 1g).

Actin oxidation by MICAL1 thus accelerates the disassembly of filaments by accelerating both cofilin-1 recruitment and subsequent severing. In other words, global severing is boosted by the increased severing rate per domain, and by the formation of numerous cofilin-1 domain boundaries whence severing occurs.

### MICAL1-oxidized actin filaments are efficiently disassembled by S3D-cofilin-1

*In vivo*, cofilin-1 is inhibited upon Ser3 phosphorylation by LIM kinases. To study the impact of this inhibition *in vitro*, we used the common phosphomimetic S3D-cofilin-1 (Blanchoin et al., 2000; Elam et al., 2017; Huehn et al., 2020). We first tested the mutant by exposing non-oxidized filaments to 1 µM unlabelled S3D-cofilin-1 (Figure 2b). After 90 seconds, we did not observe any filament severing event. We also tried to observe the nucleation and growth of cofilin-1 domains by exposing filaments to 1 µM eGFP-S3D-cofilin-1, but could not observe any binding. As a consequence, we could not measure the severing rate at the boundary of S3D-cofilin-1 domains. These observations are consistent with previous work that showed a reduced affinity of phosphomimetic cofilin-1 to non-oxidized actin filaments (Elam et al., 2017).

In a cellular context, it is expected that cofilin-1 is activated by a phosphatase, e.g. slingshot or chronophin (Kanellos and Frame, 2016). We hypothesized that actin oxidation by MICAL1 might be enough to trigger filament disassembly by S3D-cofilin-1. To test this idea, we exposed 10 µm-long oxidized actin filament segments to 1 µM unlabelled S3D-cofilin-1, and found that all filaments had severed at least once in less than 20 s (Figure 2b).

While S3D-cofilin-1 efficiently disassembles oxidized actin filaments, it still requires a much higher concentration than WT cofilin-1 on oxidized filaments (Figure 1c): the global filament severing rate at 1 µM S3D-cofilin-1 is comparable to that of 20 nM WT cofilin-1. In order to clarify this difference, we independently measured the growth rate of phosphomimetic cofilin domains and the severing rate per domain. We found that single eGFP-S3D-cofilin-1 domains grew 10 times slower than WT cofilin-1 domains onto oxidized filaments, at ∼10 sub/s/µM (Figure 2c). Surprisingly, domains of WT or S3D-cofilin-1 severed oxidized filaments equally fast (at 0.1 /s, Figure 2d).

Overall, actin oxidation bypasses cofilin-1 activation by allowing S3D-cofilin-1 to bind actin filaments, and by making filaments vulnerable to cofilin-induced severing.

### MICAL1 can oxidize Tpm1.8-decorated filaments and does not change Tpm1.8 affinity

In mammals, more than 40 Tpm isoforms are expressed and decorate the majority of actin filaments (Meiring et al., 2018). Each isoform regulates the recruitment and activity of specific ABPs (Gateva et al., 2017; Gunning and Hardeman, 2017; Manstein et al., 2019). Since the activity of MICAL1 has never been tested *in vitro* on Tpm-saturated filaments, we first wondered whether Tpm could prevent actin oxidation.

To study this question we selected a cytosolic, low molecular weight tropomyosin isoform, Tpm1.8. Brayford et al. showed that Tpm1.8 regulates actin filaments turn-over *in vivo* and promotes the persistence of the lamellipodia (Brayford et al., 2016; Gateva et al., 2017; Jansen and Goode, 2019). We recently characterized the dynamics of Tpm1.8 domains. We found that Tpm1.8 saturates the two strands of the actin filaments independently and that the domain assembly and disassembly is asymmetric: faster at the domain border closer to the filament barbed end (Bareja et al., in revision).

To detect the oxidation of filaments in our microfluidic setup, we pre-saturated actin filaments with neonGreen-Tpm1.8, and measured the barbed end depolymerization rate in the absence of G-actin in solution, in the presence or absence of MICAL1. Of note, since Tpm hardly binds to fluorescently labelled actin filaments (Gateva et al., 2017; Gunning and Hardeman, 2017; Manstein et al., 2019) we used unlabelled actin in all experiments with Tpm1.8. Previous studies on bare actin filaments showed that oxidation accelerates barbed end depolymerization about 10-fold, from ∼8 sub/s to ∼80 sub/s (Frémont et al., 2017a; Grintsevich et al., 2017).

We first measured the depolymerization rate of Tpm1.8-saturated ADP-actin filaments in the absence of MICAL1. We found Tpm-decorated filaments to depolymerize steadily at a slower rate than bare actin filaments (1.8 sub/s vs 8 sub/s, Figure 3a, b). Tpm1.8 thus appears to have a stabilizing effect, in the absence of other ABPs.

We then continuously exposed the Tpm-decorated filaments to MICAL1, from time t=0 onwards. We found that the depolymerization rate increased over time to reach a plateau around 30 sub/s (Figure 3a, b), which is still slower than the barbed end depolymerization of bare oxidized filaments (Frémont et al., 2017a; Grintsevich et al., 2017). We thus concluded that MICAL1 can indeed oxidize Tpm-decorated filaments and that the oxidation accelerates the barbed end depolymerization rate 15-fold.

Once filaments have been oxidized, we wondered whether Tpm1.8 domains would be destabilized. We thus measured the disassembly rates of neonGreen-Tpm1.8 domains in the presence of G-actin only (to prevent filament depolymerization). As was reported previously, Tpm1.8 disassembly is asymmetric (Bareja et al., in revision), and we only compared the faster and thus more accurate disassembly at the barbed end side of the Tpm domains. We found that Tpm domains disassembled as fast from non-oxidized as oxidized filaments (Figure 3c, d). Thus oxidation by MICAL1 has apparently no impact on Tpm affinity.

### Tpm1.8 protects non-oxidized filaments from cofilin-1 binding and filament severing

Many Tpm isoforms have been shown to inhibit cofilin-1 binding (Gateva et al., 2017; Jansen and Goode, 2019). In lamellipodia, Tpm1.8 has been shown to regulate filament stability along with cofilin-1 (Brayford et al., 2016) but their cooperation or competition has never been tested *in vitro*. We first measured the impact of Tpm1.8 on the activity of cofilin-1 in the absence of MICAL1.

We first observed that some cofilin-1 domains could still assemble on Tpm-saturated filaments, and we confirmed that the mCherry-cofilin-1 and neonGreen-Tpm1.8 fluorescence signals never overlap, as previously observed (Christensen et al., 2017; Gateva et al., 2017; Jansen and Goode, 2019): cofilin-1 binding induces the unbinding of Tpm. However, the majority of cofilin-1 domains nucleate from the free barbed end (Figure 4a, central kymograph). A single Tpm1.8 dimer binds along the length of filaments to six actin subunits. The barbed end most likely contains a few bare actin subunits whence cofilin-1 domains nucleate. We also observed a dramatic decrease in the nucleation rate of cofilin domains within the Tpm-decorated filament: while bare filaments get fully saturated by cofilin-1 in about 10s (at 1 µM mCherry-cofilin-1, Figure 4a left panel), more than 60% of 5 µm-long Tpm-decorated actin segments are still void of any cofilin-1 domain after 180 seconds (Figure 4b).

We then measured the growth rate of single cofilin-1 domains on bare vs. Tpm-decorated filaments. Surprisingly, we found the growth rate of cofilin domains to be reduced by only 30% on Tpm1.8-saturated filaments, compared to bare filaments (∼20 sub/s/µM on bare vs ∼13 sub/s/µM on Tpm-decorated filaments, Figure 4c left). We also measured that the growth rate was barely reduced by the addition of 50 to 100 nM Tpm1.8 in solution (Figure 4c left). We note that cofilin-1 domains could grow, and thus replace Tpm molecules on the filament, faster than the Tpm off-rate we measured in the absence of cofilin (Figure 3d). This observation indicates that cofilin does not simply replace departing Tpm molecules: cofilin accelerates the detachment of Tpm at domain boundaries. Overall, Tpm domains prevent the nucleation of cofilin-1 domains but they moderately slow down the growth of already established cofilin-1 domains.

Cofilin-1-induced severing occurs at the boundary of cofilin domains, we wondered if the presence of a contiguous Tpm domain might affect the severing rate. We discovered that the severing rate per cofilin-1 domains was reduced by 6-fold on Tpm-saturated filaments (from 2×10^−3^ /s to 3×10^−4^ /s, Figure 4d). Our results show that Tpm1.8 has an impact on every step of cofilin-1-induced disassembly, and thereby very efficiently maintains filaments.

### Oxidation allows the rapid decoration and severing by cofilin-1 of Tpm1.8-saturated filaments

Since MICAL1 can oxidize Tpm1.8-saturated filaments (Figure 3), we tested whether actin oxidation could alter the competition between Tpm and cofilin-1. We thus polymerized actin filaments and aged them in the presence of both MICAL1 and Tpm. These oxidized, Tpm-saturated filaments were then exposed to 0.1 to 1 µM mCherry-cofilin-1.

We first observed that cofilin-1 domains nucleated very rapidly over the whole filament: virtually all 5 µm-long oxidized actin segments had at least one visible cofilin-1 domain in less than 5 s (1 µM mCherry-cofilin-1, Figure 4a right panel). Oxidation thus increases the nucleation rate on Tpm-saturated filaments by ∼500-fold (from 1×10^−6^ /s/sub to 6×10^−4^ /s/sub, Figure 4b).

We then measured the normalized growth rate of cofilin-1 domains and similarly found an increase by 5-fold (from ∼13 sub/s/µM to ∼65 sub/s/µM) after oxidation of Tpm-saturated filaments. We noted that the fluorescence signal of neonGreen-Tpm1.8 and mCherry-cofilin-1 did not overlap on oxidized filament either, indicating that cofilin-1 binding leads to the very quick unbinding of Tpm. Overall, Tpm1.8 decoration does not prevent the rapid binding of cofilin-1 to oxidized filaments.

Finally we tested whether Tpm could inhibit the severing of oxidized filaments by cofilin-1. We found that single cofilin-1 domain severed Tpm-decorated oxidized filaments very rapidly, with a half-life of about 15s (Figure 4e). Therefore, oxidation by MICAL1 increases the severing rate per cofilin-1 domain by 150-fold, compared with non-oxidized Tpm-decorated filaments (from 3×10^−4^ /s to 5×10^−2^ /s). Compared to bare filaments, Tpm moderately slows down the cofilin-induced severing rate of MICAL1-oxidized filaments, by less than 2-fold (from 7×10^−2^ /s to 5×10^−2^ /s, figure 4e).

Overall, our results show that MICAL1 makes tropomyosin-decorated actin filaments extremely vulnerable to cofilin. Oxidation by MICAL1 thus appears as a powerful way to sever filaments, by upending conditions where non-oxidized filaments would be protected, without specifically removing tropomyosins and without activating large amounts of cofilin. To illustrate this idea, we sought to monitor the effect of MICAL1 activation in what would be a likely physiological situation, where filaments are decorated by tropomyosin and most of the available cofilin is phosphorylated. We thus polymerized unlabelled non-oxidized actin filaments and first exposed them to 100 nM neonGreen-Tpm1.8, and a mixture of 100 nM WT mCherry-cofilin-1 and 1 µM unlabelled S3D-cofilin-1 (Figure 5a). We observed that Tpm rapidly decorated the actin filaments, preventing cofilin-1 binding and filament severing. After 100s, the filaments were intact. We then mimicked the activation of MICAL1 by adding 500 nM MICAL1 to the solution. Over the next 100s, MICAL1 oxidized the Tpm-saturated filaments, allowing cofilin-1 to assemble into domains that rapidly fragmented all the filaments.

## DISCUSSION

We found that oxidation by MICAL1 makes filaments vulnerable to cofilin-1-induced disassembly in conditions that would otherwise be of little consequence. Trace amounts of active cofilin-1, or larger amounts of phosphomimetic S3D-cofilin-1, which are harmless to actin filaments, readily disassemble them as soon as they are oxidized by MICAL1 (Figures 1, 2, 5). Filament decoration by Tropomyosin Tpm1.8, which offers an efficient protection against cofilin-1-induced disassembly, becomes inefficient as soon as the filaments are oxidized (Figures 3, 4, 5).

Filament oxidation by MICAL1 can thus be viewed as a way to abolish different protections from cofilin-induced disassembly. Interestingly, the temporary protection offered by actin’s ADP-Pi content, which prevents cofilin from binding to freshly assembled filament regions (Suarez et al., 2011) is also suppressed by MICAL1-induced oxidation (Grintsevich et al., 2017). MICAL1-induced oxidation thus appears to favor the action of cofilin in various situations.

Remarkably, these multiple effects, which upend the regulation of actin disassembly, are due to the modification of only two residues in the D-loop of actin, Met44 and Met47 (Hung et al., 2011). As shown previously, these modifications weaken intersubunit bonds and, as a consequence, filaments depolymerize faster (Frémont et al., 2017a; Grintsevich et al., 2017) and are easier to fragment by mechanical shearing (Grintsevich et al., 2016). Here, our experiments apply negligible mechanical stress to the filaments, and we further confirm that oxidized filaments barely sever without cofilin (Figure 1). As with non-oxidized filaments, severing occurs at the boundaries of cofilin domains and preferably at the pointed end boundary of cofilin domains. These observations suggest that filament oxidation does not alter the nature of the severing mechanism, and that the weakening of intersubunit bonds may suffice to explain the spectacular 50-fold increase in the severing rate at cofilin domain boundaries (Figure 1g).

In addition, the oxidation of the D-loop of actin favors the binding of cofilin, and especially the binding of the first cofilin molecules (Figure 1e), leading to the formation of many cofilin domains and domain boundaries, thereby further increasing the occurrence of severing events. Structural data show that, when cofilin binds to an actin filament, the N-terminal region of cofilin contacts the vicinity of actin’s D-loop (Blanchoin et al., 2000; Elam et al., 2017; Galkin et al., 2011; Huehn et al., 2020). Local changes induced by oxidation in the D-loop (Grintsevich et al., 2017) may thus explain how cofilin binding is favored. Further, this local interaction is known to be impaired by modifications on Ser3 of cofilin (Huehn et al., 2020) and our results suggest that it may be restored by the oxidation of the D-loop of actin (Figure 2). The oxidation of actin’s D-loop may also induce global modifications of the filament conformation and stiffness, which have yet to be measured, and which could contribute to favor cofilin binding.

It appears that most Tpm isoforms inhibit cofilin-induced disassembly of non-oxidized filaments, to different extents (Gateva et al., 2017; Jansen and Goode, 2019). Here, we show that Tpm1.8 can efficiently prevent the nucleation of new cofilin domains and that, in addition, it strongly reduces severing at cofilin-Tpm domain boundaries (Figure 4). Tpm binding and MICAL1-induced oxidation appear unaffected by each other (Figure 3), consistent with the observation that Tpm binds actin filaments along their positively charged groove, away from the D-loop (von der Ecken et al., 2015). Nonetheless, Tpm1.8 prevents the formation of cofilin domains on non-oxidized actin filaments, and is no longer able to do so when the D-loop is oxidized (Figure 4). It is thus likely that more global effects, such as perhaps changes in filament stiffness induced by Tpm binding (Greenberg et al., 2008), also play a role and need to be explored further.

Our results illustrate how factors of different natures can come together to regulate actin assembly (Jégou and Romet-Lemonne, 2016). The interplay between ABPs and mechanical factors, in particular, is currently under intense scrutiny (Harris et al., 2018; Jegou and Romet-Lemonne, 2019; Schramm et al., 2017; Suzuki et al., 2020; Wioland et al., 2019b; Zimmermann and Kovar, 2019). Today, PTMs of actin are emerging as a key regulatory factor of actin assembly (Varland et al., 2019) and how they modify interactions with ABPs is mostly unknown. Here, we have quantified the role of the direct PTM of actin by oxidation as a critical regulatory mechanism that controls actin disassembly, in synergy with well-established ABPs. In the future, we expect that more studies will go beyond the simple combination of ABPs, to decipher the multifactorial regulation of the cytoskeleton.

We propose that, in cells, the local activation of MICAL1 is a powerful way to trigger the severing of actin filaments, by turning innocuous concentrations of active and inactive cofilins into a lethal mix for filaments, and by making tropomyosins powerless to protect the filaments. Several ABPs are known to assist cofilin to rapidly disassemble filaments, and they may also be sensitive to the change in the redox state of actin filaments.

## ACKNOWLEDGEMENTS

This work was supported by the CNRS, Institut Pasteur, the ANR (grant Cytosign to A.E, and grant RedoxActin to A.E. and G.R.L.) and the ERC (grant StG-679116 to A.J.). The authors thank Till Boecking for the plasmid of Tpm1.8.

## METHODS

### Proteins purification and labelling

Skeletal muscle actin was purified from rabbit muscle acetone powder following the protocol described in (Wioland et al., 2017), adapted from the original protocol (Spudich and Watt, 1971). Actin was labelled on accessible surface lysines of F-actin, with Alexa-488 or Alexa-568 succinimidyl ester (Life Technologies). To limit artifacts from the fluorophore, we used a labelling fraction of 12 %, except in experiments with Tpm where filaments were unlabelled.

Spectrin-actin seeds were purified from human erythrocytes as described (Wioland et al., 2017), based on the original protocol by (Casella et al., 1986).

Mouse cofilin-1 (Uniprot : P18760) and S3D-cofilin1, unlabelled or fluorescently tagged with eGFP and mCherry at their N-terminus, were purified as described previously in (Kremneva et al., 2014).

neonGreen-Tpm1.8 was constructed as a protein with N-terminal His6-tag followed by mNeonGreen and a peptide linker (GGGSGGGSGTAS) fused to the N-terminus of Tpm1.8, as described in (Bareja et al, in revision). The alanine-serine fused directly at the N-terminus of Tpm1.8 mimics its acetylation.

Since the full length MICAL1 is auto-inhibited, we only used the catalytic domain of human MICAL1, purified as described in (Frémont et al., 2017a). It was always supplemented with its cofactor nicotinamide adenine dinucleotide phosphate (NADPH, 12 µM).

### Buffer

We performed all experiments in standard F-buffer (5 mM Tris HCl pH 7.4, 50 mM KCl, 1 mM MgCl_2_,0.2 mM EGTA, 0.2 mM ATP, 10 mM DTT and 1 mM DABCO).

### Microfluidics chamber

Experiments were performed in Poly Dimethyl Siloxane (PDMS, Sylgard) microfluidics chambers, 20 µm in height, 800 µm in width and about 1 cm long. Chambers had a cross-shape with 3 inlets, merging into a central channel, leading to the outlet (Jégou et al., 2011).

Coverslips were washed and sonicated with Helmanex (30 min sonication), 1M KOH (30 min) and pure ethanol (30 min) and thoroughly rinsed with dH20 between the three steps. Coverslips were finally dried with pressurized air and exposed to UV along with PDMS chambers to allow for a tight binding.

### Chamber functionalization and passivation

Twenty µL of the following solutions were directly injected into the chamber with a pipette. First 18 pM Spectrin-actin seeds, left for 30 to 60s for seeds to adhere non-specifically to the glass surface. Followed by PLL-PEG (1 mg/mL, 1h), casein (15 min) and BSA (15 min) to passivate the surface.

### Filament elongation and experiment

Experiments were performed in three steps (Figure 1a):

1. Filament elongation: grown from seeds with 0.6 to 1 µM ATP-G-actin for about 10 min. Filaments are anchored to the surface by their pointed end only, through a spectrin-actin seed.
2. Filament elongation, oxidation and Tpm-saturation: filaments are then aged for 15 minutes with a solution of ATP-G-actin at critical concentration, ∼0.1 µM. This results in more than 99.9% of the subunits in an ADP form. Actin subunits can also be oxidized at this step by supplementing the solution with typically 50 nM FAD and 12 µM NADPH. Simultaneously filaments can be saturated with 100 nM Tpm1.8.
3. Filament or Tpm domain disassembly: filaments are finally exposed to buffer, in the presence or absence of WT or S3D-cofilin-1.

### Acquisition

The microfluidic setup was placed on a Nikon TiE or TE2000 inverted microscope, equipped with a 60x oil-immersion objective and an optional additional 1.5x magnification. We either usedTIRF, HiLo or epifluorescence depending on the background fluorophore concentration in solution. The TiE microscope was controlled by Metamorph, illuminated in TIRF or epifluoresence by 100 mW tunable lasers (ILAS2, Roper Scientific), and images were acquired by an Evolve EMCCD camera (Photometrics). The TE2000 microscope was controlled by micromanager (Edelstein et al, 2014), illuminated with a 120W Xcite lamp (Lumen dynamics) and images were acquired by an sCMOS Orca-Flash2.8 camera (Hamamatsu).

### Analysis

All experiments were performed at least twice and at least one representative movie was analyzed with the following procedures:

- “Filament severing”: a set of randomly picked 10 µm-long actin segments was monitored from time t=0 when cofilin (or buffer) was first injected. Segments will either sever or be lost due for example to severing somewhere else on the filament (e.g. at the anchoring). We calculated the survival fraction from this set of severing or loss times, using a standard Kaplan-Meier method. Survival fraction, confidence intervals and p-values (log-rank test) were calculated using the Python package Lifelines. Survival fractions were fitted with a single exponential on Excel (Microsoft Office) to obtain reaction rates (shown as thin continuous grey lines on plots).
- “Severing per cofilin domain”: we tracked single cofilin domains, from the frame on which they appear (nucleation) which defines t=0. We then followed the same procedure as for “filament severing”.
- “Cofilin domain nucleation”: similarly, 5 µm long actin segments were tracked from t=0 when cofilin is injected. When then measured the frame on which a cofilin domain nucleates or the segment is lost. The survival fraction was calculated with the same Kaplan-Meier method.
- “Cofilin domain growth”: the growth rate was measured by two different methods. When large domains can assemble (i.e. when nucleation is limited), the growth rate is simply the sum of the rates towards both ends, measured manually on kymographs made with ImageJ. When many domains nucleate simultaneously and rapidly merge, we measure the fluorescence intensity of a single domain spreading over maximum 3 pixels. The intensity was normalized by that on a filament fully saturated with cofilin. We previously showed that the two methods yield comparable results (Wioland et al, 2017). P-values are calculated with a Student’s t-test.
- “Tpm disassembly”: similarly we measured the disassembly rate of Tpm domains directly on kymographs. However we only focused on the disassembly towards the pointed end which has been shown to be faster (Bareja et al., in revision) and thus more accurate.
- “Barbed end depolymerization”: filament barbed end position was measured on every frame with a custom build algorithm on Python. The mean depolymerization rate at a time t was obtained by pooling the barbed end position of all filaments over 5 frames (centered around time t). The position over time was fitted with a linear function whose slope yields the depolymerization rate.

---

**Supp. Fig. 1.**
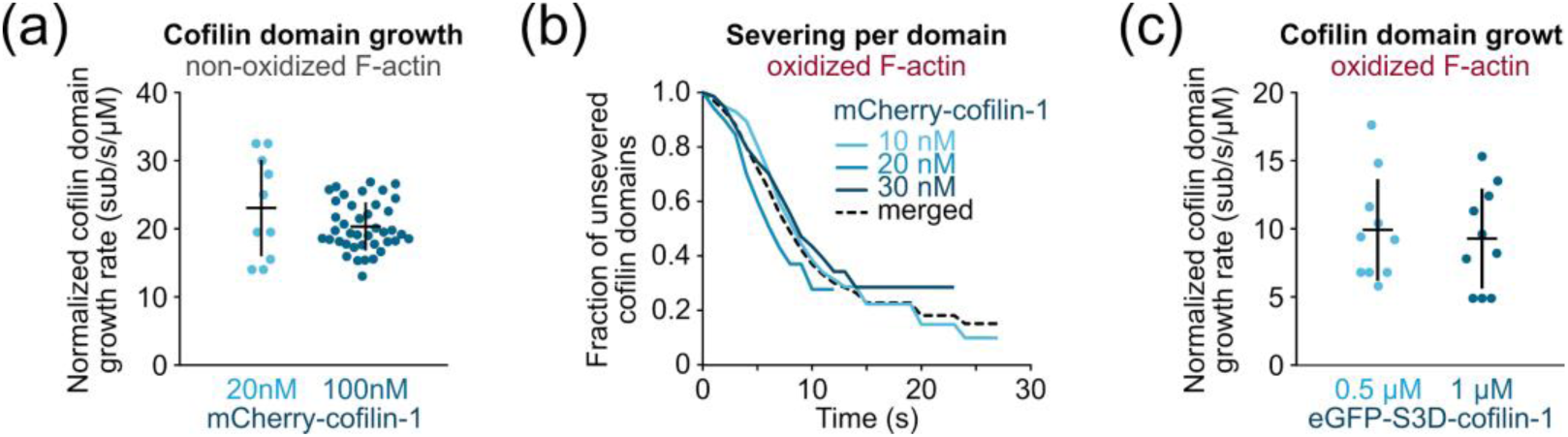
The normalized cofilin domain growth rate and the severing rate per cofilin domain are independent of cofilin concentration. (a) growth rate of WT mCherry-cofilin-1 domains on non-oxidized actin. N = 10 domains (20 nM cofilin) and 40 domains (100 nM cofilin). Bars: mean and S.D. The data were pooled together for Figure 1f. (b) Filament severing at single cofilin-1domains. N = 60 (10 nM cofilin), 70 (20 nM cofilin) and 75 domains (30 nM cofilin). The data were pooled together for Figure 1g. (c) Growth rate of eGFP-S3D-cofilin-1 domains on oxidized actin. N = 10 domains for each condition. Bars: mean and S.D. The data were pooled together for Figure 2c.

